# Nanoscale visualization of ribonucleoprotein condensates with ultrastructure RNA expansion microscopy

**DOI:** 10.64898/2026.06.02.729447

**Authors:** Jeet Mukhopadhyay, Parakbritta Hazra Dutta, Samiran Bhowmik, Pintu Patra, Mainak Bose

## Abstract

Superresolution imaging of RNAs in their native cellular context is crucial for investigating RNA biology. Here, we develop Ultrastructure RNA Expansion (UREx) microscopy, an expansion microscopy approach, that retains cellular RNAs and RNP granule condensates. We demonstrate that UREx retains the highly ordered core-shell architecture of *Neat1* RNA in the paraspeckles, confirming RNA ultrastructure preservation. In addition to robust RNA retention, UREx also preserves protein epitopes and overall cellular protein ultrastructure thus enabling context-dependent visualization of RNAs in cells as well as intact tissues. Employing UREx, we visualize RNP condensates in the intact *Drosophila* germline and resolve diffraction-limited *oskar* granules into RNP nanoclusters. Together, our findings establish UREx as a broadly applicable and cost-effective resource to interrogate the nanoscale spatial organization of RNAs and proteins within diffraction-limited RNP assemblies.

## Introduction

The genome codes for a vast number of RNAs, of which messenger RNAs (mRNAs) are translated into proteins while others are non-coding in nature, including ribosomal RNAs (rRNAs), transfer RNAs (tRNAs), long non-coding RNAs (lncRNAs), etc. Throughout their lifespan, both mRNAs and non-protein coding RNAs dynamically associate with RNA-binding proteins (RBPs), forming ribonucleoprotein (RNP) complexes and undergo regulated processing, transport, translation, and degradation to maintain cellular homeostasis. Within the crowded intracellular environment, RNAs are often not randomly dispersed but show specific localization patterns, can be targeted to organelles or stored/sequestered in membraneless RNP granules^1^. Although transcript abundance can be quantified by PCR or sequencing-based approaches, these methods lack the information about the spatial context which is circumvented by RNA visualization with light microscopy. Therefore, high-resolution imaging of RNAs at various stages of their complex life and in their subcellular location is key to understanding how gene expression is spatially organized.

Nucleic acid hybridization-based methods, such as single molecule Fluorescence In Situ Hybridization (smFISH), aid visualization of single transcripts in fixed specimens but questions related to nanoscale RNA organization cannot be addressed using conventional diffraction-limited microscopy. Confocal microscopy restricts the spatial resolution to ∼200–250 nm precluding visualization of cellular structures close to this limit. However, RNPs can exist and function at length scales below the diffraction limit. Therefore, internal architecture and spatial organization within diffraction-limited assemblies such as, RNP granules, cannot be resolved using conventional microscopy. The advent of super-resolution microscopy circumvents this problem by surpassing the diffraction limit employing optical as well as computational strategies. Stimulated emission depletion (STED) microscopy reduces the effective point spread function (PSF) by selectively depleting fluorescence around the excitation focal spot to achieve sub-diffraction resolution^2,3^. On the other hand, single-molecule localization microscopy (SMLM) techniques rely on the stochastic activation and precise localization of individual fluorophores over time to reconstruct images with nanometer resolution^4^. However, complex sample preparation, harsh imaging conditions and the requirement of specific dyes limit their applications, especially for complex specimen and intact tissue samples.

The advent of Expansion Microscopy (ExM) has provided a complementary route to super-resolution imaging by physically magnifying the specimen instead of optically overcoming the diffraction limit. In ExM, the biomolecules in the sample are anchored to a swellable polymer network, which can be isotropically expanded through hydration^5^. This increase in physical distances between biomolecules that were originally below the diffraction limit permits super-resolution imaging using conventional microscopes. Combining FISH methodologies with ExM has aided in nanoscale visualization of RNAs in cell monolayers and thin tissue sections. These methods either rely on covalently anchoring RNA molecules (via Guanine bases) to the acrylamide (AA) gel or using acrydite-modified FISH probes^6,7^. However, protein epitopes and the overall cellular ultrastructure are not preserved in these approaches, thereby precluding visualization of RNAs in association with RBPs and in their native cellular context. Moreover, the use of specific chemicals for anchoring RNAs or customized probes limits their routine laboratory application.

Recently, it has been shown that Ultrastructure expansion microscopy (U-ExM) preserves cellular ultrastructure and molecular architecture of subcellular complexes^8^. U-ExM is a robust expansion microscopy protocol that is extensively employed due to its ability to preserve nanoscale cellular architecture with high-fidelity, as benchmarked by the preservation of the nine-fold microtubule triplet symmetry in centrioles^8^. U-ExM uses a combination of acrylamide (AA) and formaldehyde (FA) to anchor biomolecules post-fixation, followed by hydrogel embedding and in situ polymerization. The classical pro-ExM protocol employs proteolytic digestion to homogenize the specimen thus allowing isotropic expansion of the gel upon addition of water^1^. In contrast, U-ExM samples are homogenized by heating with detergent (SDS) at elevated temperature and this controlled denaturation also facilitates enhanced ultrastructure preservation. However, it remains unclear if RNAs are retained and can be imaged using this method. While recently developed expansion microscopy approaches such as, Click-ExM^9^ and Magnify^10^ have introduced robust biomolecule anchoring strategies capable of retaining proteins, lipids, nucleic acids in the gel matrix, direct demonstration of RNA ultrastructure preservation is missing.

Here, we develop Ultrastructure RNA Expansion microscopy (UREx), an extension of U-ExM, that improves RNA retention in the gel and aids in super-resolution imaging of RNAs and RNP granules in cells. Importantly, UREx preserves protein epitopes and overall cellular ultrastructure thus allowing probing RNAs in their native subcellular context by conventional in situ hybridization techniques.

## Results

### Ultrastructure RNA Expansion (UREx) microscopy preserves cellular RNAs

In the anchoring step, U-ExM uses a mixture of 0.7% FA and 1% AA to covalently link cellular proteins to the polymer matrix^11^. FA reacts with primary amines of lysine residues in proteins to generate methylene bridge crosslinks with concomitant incorporation of AA moieties between biomolecules. Thus, biomolecules become chemically functionalized with AA monomers and during gelation are covalently integrated into the polyacrylamide-based hydrogel matrix. Since RBPs coat cellular RNAs, which during fixation gets covalently crosslinked to the bound proteins, we reasoned that the conventional U-ExM protocol should retain cellular RNAs. However, performing U-ExM on paraformaldehyde (PFA)-fixed CHO cells, followed by 28S rRNA smFISH, we observed that 28S rRNA was barely detectable even at a ten-fold higher smFISH probe concentration relative to unexpanded controls (Extended Data Fig. 1A). Although the expected localization of 28S rRNA to the cytoplasm and the nucleolus was retained, the weak signal suggested inefficient retention of RNA within the expanded gel matrix.

Proteins are present at substantially higher abundance than RNA in mammalian cells, comprising ∼18–20% of total cellular mass compared to ∼1% for RNA^12^. Owing to their higher abundance, both in terms of mass and copy number, protein ultrastructure is more efficiently retained in U-ExM. RNA is primarily stabilized through RNA–protein and protein–protein crosslinking networks, which immobilize transcripts and preserve their spatial position. To improve direct anchoring of RNAs via their primary amines (Adenine C_6_, Guanine C_2_, Cytosine C_4_), we increased the FA concentration in the anchoring solution. We observed a significant 3-fold enhancement of 28S rRNA retention, as compared to U-ExM, with our anchoring solution containing 2.1% FA + 1% AA (Extended Data Fig. 1A-B). Notably, no significant effect on the expansion factor was observed (Extended Data Fig. 1C). We term this procedure as Ultrastructure RNA Expansion (UREx) microscopy (Fig. 1A). Employing the UREx workflow, we simultaneously visualized the subcellular distribution of rRNA and polyadenylated mRNA using 28S rRNA and oligo(dT) smFISH probes, respectively, in cultured CHO cells (Extended Data Fig. 2).

**Fig. 1:**
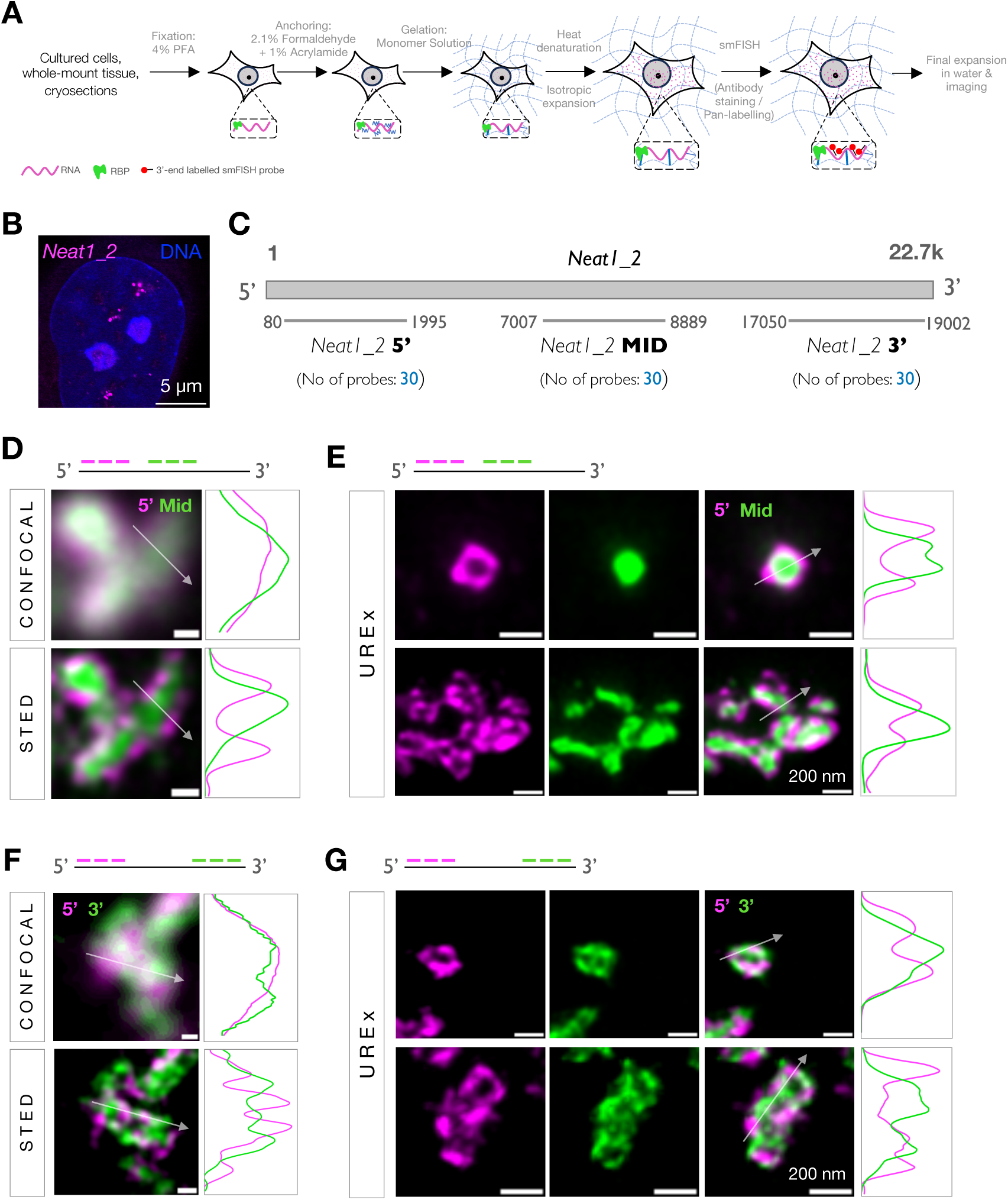
Ultrastructure RNA Expansion (UREx) microscopy preserves core-shell organization of *Neat1* in paraspeckles. (A) Schematic workflow of the UREx protocol. (B-C) Paraspeckles detected by smFISH against *Neat1_2* RNA in Hela cells (B). Schematic representation of human *Neat1_2* RNA indicating the regions against which smFISH probes are designed. (D-E) Representative confocal and STED images of paraspeckles (D) probed against the 5’ (Atto 633, magenta) and Mid (Star 580, green) regions of *Neat1_2*. Corresponding UREx images (E) show a representative spheroidal and oblate/sausage-like paraspeckles. For UREx, smFISH probes against 5’ and Mid region were labelled with Atto 633 and Atto 565, respectively. (F-G) Representative confocal and STED images of paraspeckles probed against the 5’ (Atto 633, magenta) and 3’ (Star 580, green) regions of *Neat1_2*. Corresponding UREx images are shown in G. For UREx, smFISH probes against 5’ and 3’ regions were labelled with Atto 633 and Atto 565, respectively. Normalized intensity line profiles (grey arrow) spanning 1 µm (real space) along the paraspeckle are depicted in panels D-G. In all expansion images, the scale bars are rescaled according to the expansion factor.

### UREx preserves spatial organization of *Neat1* in paraspeckles

Multivalent interactions between individual RNP complexes can lead to their condensation into higher-order RNP granules. A plethora of studies have now highlighted the emerging role of RNP condensation in RNA processing, localization, translation, stress response, etc^13^. Recent evidence suggests that RNAs can play an architectural role in scaffolding condensate assembly^14^. One such RNA is *Neat1* lncRNA which forms the architectural backbone of paraspeckles. Paraspeckles are sub-micron nuclear condensates scaffolded by the long isoform of *Neat 1*, known as *Neat1_2*, a 22.7kb long transcript in humans^15,16^ (Fig. 1B). In addition, several paraspeckle proteins (PSPs), such as, NONO, SFPQ, FUS, etc accumulate in these nuclear bodies^17^. Depletion of *Neat1_2* isoform leads to disassembly of paraspeckles, confirming the structural role of the lncRNA in paraspeckle assembly^18^. Using electron microscopy (EM) and structured illumination microscopy (SIM), it has been shown that *Neat1_2* RNA is radially organized with the 5’ and 3’ ends projecting towards the condensate periphery and the middle domain at the centre of paraspeckles^19–21^. This results in a characteristic ‘core-shell’ architecture which is the hallmark of these condensates^22^. ExM approaches have exploited highly ordered protein assemblies, such as, centrioles and nuclear pore complexes (NPCs) to benchmark structural preservation, but analogous symmetrical RNA assemblies are rare^11,23^. To this end, we decided to leverage the well-defined spatial organization of *Neat1_2* in paraspeckles to benchmark UREx.

To assess if UREx preserves *Neat1_2* orientation within paraspeckles, we designed smFISH probes against the 5’, 3’ and mid domains of *Neat1_2* (Fig. 1C). However, we observed that fixation with 4% PFA alone was not sufficient for retention of *Neat1* in the gel matrix, as evident from the smFISH signal-to-noise ratio (Extended Data Fig. 3). Although glutaraldehyde (GA) is less optimal than PFA as a fixative for confocal microscopy due to strong autofluorescence, GA promote rigid crosslinking and superior structural preservation^24,25^. To minimize the caveats of GA fixation, we fixed the cells with very low concentration of GA (0.1%) in combination with 3% PFA. This substantially improved *Neat1* signal in the UREx expanded cell samples (Extended Data Fig. 3). Next, we performed 2-color smFISH experiments to simultaneously detect the 5’-mid or the 5’-3’ regions of the RNA in Hela cells. While conventional confocal microscopy failed to spatially resolve the 5′ and mid regions, STED imaging clearly separated these signals as evident from the intensity line profiles (Fig. 1D). The clear separation of the two signals was also evident with UREx revealing the characteristic core–shell organization of *Neat1_2* within paraspeckles (Fig. 1E). Similarly, simultaneous probing of the two ends of the RNA with UREx revealed that the 5′ and 3′ domains are arranged adjacent to each other in a complementary manner along the periphery of the paraspeckle, as also observed with STED imaging on the unexpanded samples (Fig. 1F-G). This indicates that UREx not only resolves the *Neat1_2* RNA domains distinctly but also preserves the native orientation of this architectural RNA with high fidelity.

We further performed 3-color smFISH with UREx and simultaneously visualized the spatial organization of the 5’, mid and 3’ domains of *Neat1_2* in these condensates (Fig. 2A). High-resolution light microscopy as well as electron microscopy studies have established that paraspeckles are cylindrical bodies usually detected as spheroids (transverse plane) or as irregular sausage-like structures (longitudinal plane) in the nucleoplasm^19^ (Fig. 1 D-G, 2A). To get a quantitative estimate of core–shell spatial organization and benchmark ultrastructure preservation with UREx, we segmented the three domains and quantified the pairwise nearest-neighbour distances between the domains. Our analysis revealed a mean separation of 52.72 ± 28 nm between the 5′ and 3′ domains of *Neat1_2*. Similarly, the distances between the middle domain and the 5′ end, and between the middle domain and the 3′ end, were 97.28 ± 57 nm and 105.3 ± 53 nm, respectively (Fig. 2B-C). The values are in agreement with the published distance for mouse *Neat1_2* calculated using SIM imaging^20^. Since paraspeckles appear as either spheroidal or cylindrical structures, centroid-based measurements might not accurately reflect the spatial proximity between the core and shell domains. Therefore, we additionally quantified the Euclidean distance between the centroid of 5’ (shell) signal to the nearest edge of the mid (core) signal, yielding a mean separation of 22.3 ± 14.2 nm (Fig. 2D).

**Fig. 2:**
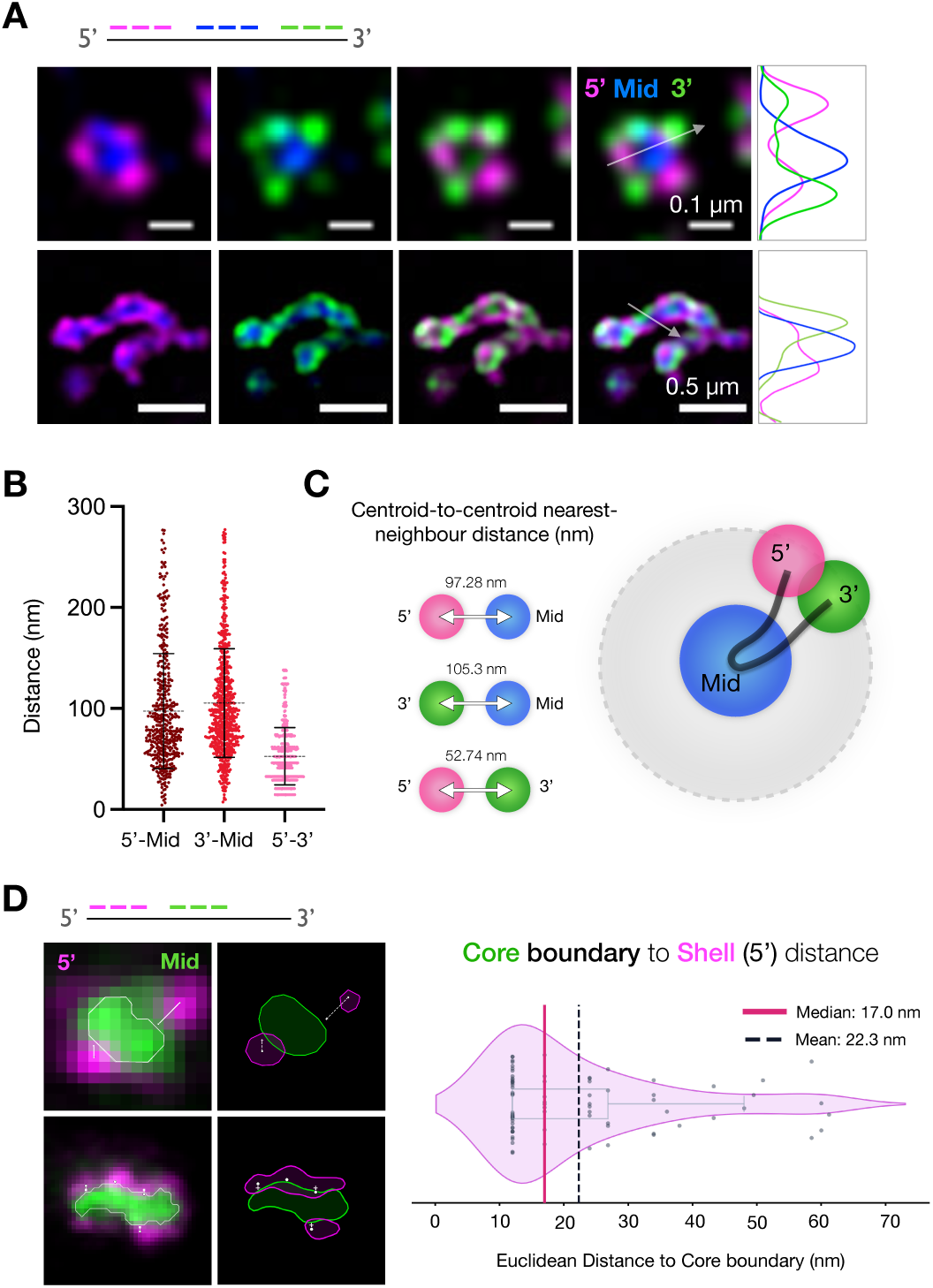
Benchmarking UREx with *Neat1* organization in paraspeckles. (A) 3-color *Neat1_2* smFISH on UREx expanded HeLa cells with probes labelling the 5’ (Atto 633, magenta), mid (Atto 488, blue) and 3’ (Atto 565, green) regions. Normalized intensity line profile (grey arrow) along the paraspeckle is depicted. (B-C) Quantification of the nearest neighbour centroid-to-centroid distances between the individual domains from multiple paraspeckles from a total of 5 FOVs (B). Model of *Neat1_2* architecture within a paraspeckle with the 5’, mid and 3’ domains and the mean nearest-neighbour distance between them indicated (C). (D) Representative 2-color smFISH UREx images of (spheroidal and elongated) paraspeckles are shown with the segmentation of the core signal (white outline) and the distance from core boundary to shell (white line). Violin plot (right) represents the quantification of the distance from core (green) surface to the centroid of the nearest 5’ (magenta) signal from multiple paraspeckles. In the expansion images, the scale bars are rescaled according to the expansion factor.

### UREx retains native cellular ultrastructure and specific protein epitopes

To test if UREx could preserve the cellular ultrastructure like U-ExM, we performed a pan-proteome labelling using dye-coupled NHS ester^26^. By reacting with primary amines, NHS esters enable dense labelling of protein-rich structures in expanded samples with high contrast. Combining NHS staining with UREx, we confirmed that nanoscale cellular architecture was preserved as evident from the labelling of NPCs, chromatin domains, mitochondria (Extended Data Fig. 4A). We next examined whether specific protein epitopes are retained following UREx and accessible to antibody labeling. Immunofluorescence (IF) was carried out using either directly tagged primary antibody or primary-secondary combination. The microtubule cytoskeleton was robustly visualized with anti-tubulin immunostaining, while anti-Tomm20 specifically stained the outer mitochondrial membrane with clear exclusion from the mitochondrial lumen (Extended Data Fig. 4B–C). Together, these results demonstrate that, similar to conventional U-ExM, UREx preserves both cellular ultrastructure as well as specific protein epitopes.

### Resolving the luminal distribution of mitochondrial transcripts with UREx

Most membrane-bound organelles including mitochondria, have dimensions close to the diffraction limit of conventional light microscopes. The mitochondrial DNA (mtDNA) is transcribed into polycistronic precursor RNAs that undergo co-transcriptional processing to generate eleven mature mRNAs, two rRNAs and several tRNAs^27^. However, the localization and spatial organization of mitochondria-encoded mRNAs remains difficult to resolve. In a recent study, Stoldt et al., combined smFISH with STED imaging to resolve three mitochondria-encoded mRNAs^28^. We set out to test if UREx enables visualization of mitochondrial RNAs with nanoscale precision.

To retain mitochondria post-expansion, Hela cells were fixed with PFA+GA as described previously for U-ExM^11^. Using UREx, mitochondria was readily detectable with mitotracker dye as well as NHS-ester staining which stained mitochondria with high-contrast. The luminal distribution of Cytochrome C oxidase subunit III (*COX3*) mRNA, interspersed with the super-resolved mitotracker signal, was detected by smFISH on the gels (Fig. 3A). Mitotrackers are cationic dyes that covalently bind to thiol-containing mitochondrial proteins. Note that, unlike in unexpanded cells, the *COX3* and the mitotracker spots are spatially resolved and seldom overlaps within the mitochondrial lumen, highlighting the resolving power of UREx (Extended Data Fig. 5). Furthermore, to quantify the resolution improvement, we segmented the mitotracker signal from unexpanded and UREx images and quantified the number of *COX3* spots per unit mitochondrial area, using Big-FISH spot detection algorithm^29,30^. In contrast to 1-2 *COX3* spots in unexpanded cells, an average of 15-20 spots per µm^2^ of mitochondria could be detected and resolved with UREx (Fig. 3B).

**Fig. 3:**
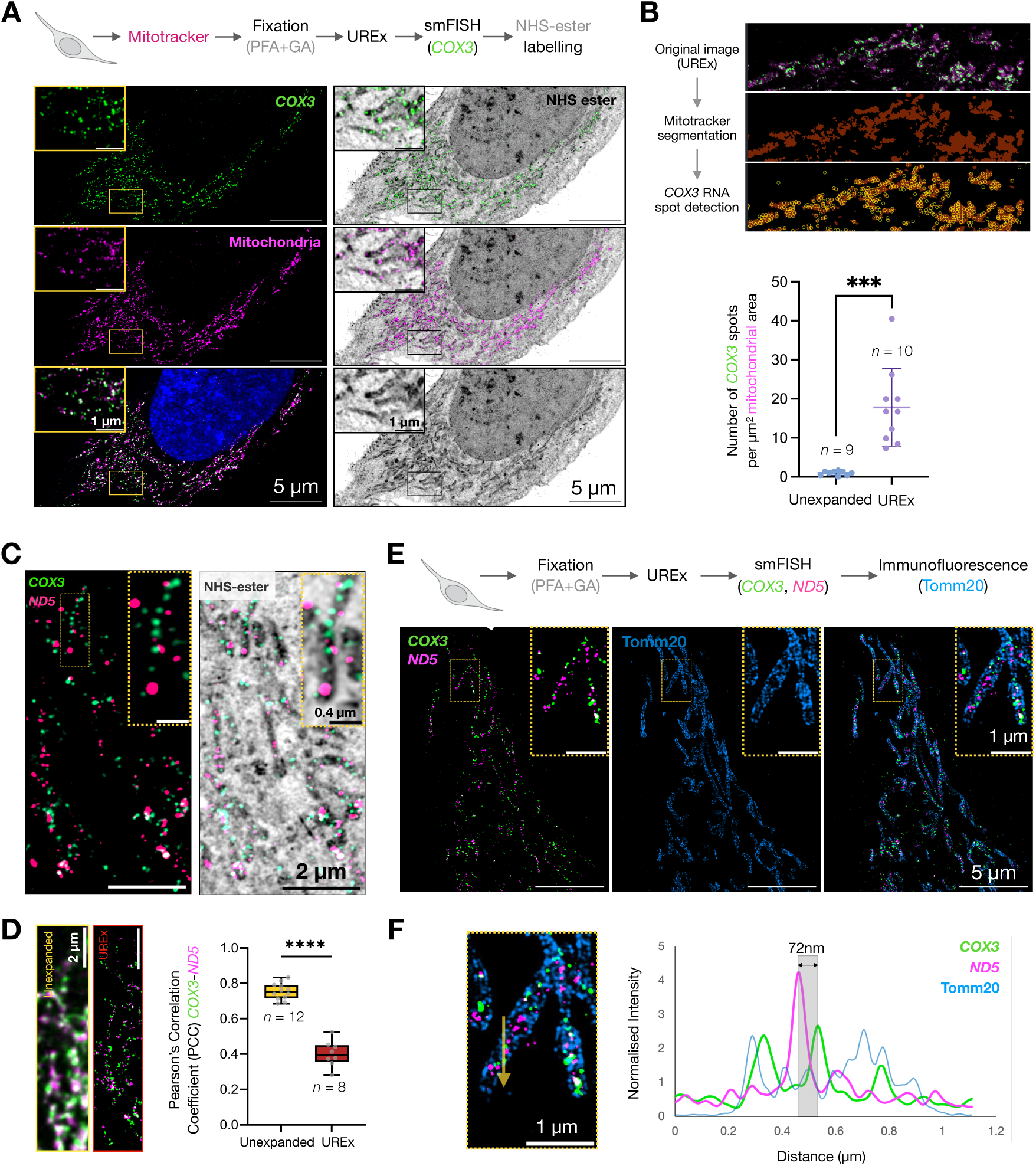
Super-resolution imaging of mitochondrial RNAs with UREx. (A) Representative confocal image of *COX3* mRNA (Atto 565) distribution on mitochondria in UREx-expanded HeLa cells, visualised by smFISH. Mitochondria have been visualized using Mitotracker Deep Red (magenta). Pan-labelling using Atto 488 NHS-ester (grey) distinctly labels the mitochondria with high contrast. Boxed areas are enlarged in the top left. DNA is stained using Hoechst. (B) Quantification of the number of *COX3* RNA spots per unit area (1 µm^2^) of mitochondria. The UREx expanded image is shown to depict the analysis workflow. Unpaired Student’s t-test was used for comparisons. Significance level: *** p<0.001. n = number of fields of view (FOVs). (C) Representative confocal image of *COX3* (Atto 565, green) and *ND5* (Atto 633, magenta) RNAs in UREx-expanded HeLa cells, visualised by smFISH. Cellular ultrastructure is visualized with Atto 488 NHS-ester pan-labeling (grey) which densely labels the mitochondria of the cell. The boxed area is magnified on the top right depicting the intraluminal resolution of the two mRNAs. (D) Quantification of the Pearson’s correlation coefficient (PCC-Costes), as a metric for colocalization between *COX3* and *ND5* RNAs in unexpanded and UREx samples. The drop in PCC indicates the improvement in resolution using the UREx workflow. Representative distribution patterns of the RNAs in unexpanded and expanded cells are shown on the top. Unpaired Student’s t-test was used for comparisons. Significance level: **** p<0.0001. n = number of fields of view (FOVs). (E) Combined smFISH and IF in UREx expanded Hela cells. Representative confocal image of *COX3* (green), *ND5* (magenta) mRNAs detected by smFISH and anti-Tomm20 IF (blue) to visualize the mitochondrial outer membrane. Boxed areas (yellow) are magnified on the top right. (F) Normalized intensity line profile (yellow arrow) across a single mitochondrion depicts the resolution enhancement with UREx. In all expansion images, the scale bars are rescaled according to the expansion factor.

We then simultaneously probed for *COX3* and another mitochondrial transcript, *ND5*, and observed distinct, non-overlapping sub-luminal distribution pattern of the two mRNAs (Fig. 3C). To quantitatively assess the spatial relationship between the two mitochondrial mRNAs, we calculated the Pearson’s correlation coefficient (PCC) as a measure of colocalization. In contrast to diffraction-limited imaging, which showed considerable overlap of the two RNAs, UREx showed a marked 2-fold reduction in colocalization, suggesting that the transcripts occupy spatially distinct domains within the lumen (Fig. 3D). Beyond the improvement in spatial resolution, the reduced PCC further suggests that the detected mRNAs predominantly exist as processed individual transcripts rather than unprocessed intermediates, consistent with the finding that mitochondrial mRNAs are immediately excised from the primary polycistronic transcript^28^.

### UREx can be combined with immunofluorescence

While NHS-ester co-staining provides the information on the subcellular context (Fig 3A), simultaneous visualization of RNAs of interest in combination with specific proteins is essential to understand cell biology. To combine smFISH with IF in the UREx pipeline, we performed a short IF step at room temperature post smFISH which successfully retained both the RNA and the protein signals (see Methods). By combining *COX3* and *ND5* smFISH with Tomm20 immunostaining, we outlined the outer mitochondrial membrane and resolved the spatial distribution of the two mRNAs within the mitochondrial lumen (Fig. 3E-F). This approach could be readily applied to simultaneously visualize RNA and RBP distribution within biomolecular condensates.

### UREx resolves diffraction-limited *oskar* RNP condensates in developing *Drosophila* embryos

ExM has been previously applied for high-resolution imaging of tissue sections, but not for intact tissues^5^. With UREx, we expanded intact early *Drosophila* embryos and performed confocal imaging of the whole-mount gel-embedded tissue using a long working-distance objective lens. In addition to visualizing the tissue ultrastructure with NHS-ester chemistry, we also confirmed that RNP condensates were preserved with UREx by IF against germ granule marker protein, Vasa (Extended Data Fig. 6A-B). In *Drosophila*, germ granules are nucleated by the maternal protein Oskar, which accumulates at the embryo posterior (germ plasm) by directed transport and local translation of *oskar* mRNA^31–33^,. *oskar* mRNAs package with specific RBPs into diffraction-limited transport RNP condensates in the *Drosophila* oocyte and condensation is critical for transport and translation of the mRNA^34^.

Interestingly, with *oskar* smFISH and conventional confocal imaging, we observed that the diffraction-limited *oskar* granules present in the somatic cytoplasm form larger condensates in the germ plasm of early embryos (Fig. 4A). We next tested if UREx could resolve the spatial RNA organization within these condensates. UREx of whole-mount embryos resolved these condensates into ‘agglomerates’ of multiple RNP nanoclusters (Fig. 4B). This indicates that multiple *oskar* RNP granules cluster together tightly into agglomerates at the embryo posterior. Agglomeration of *oskar* has been reported in late-stage oocytes, as observed by STED nanoscopy^35^. To verify that this clustering was not an artefact of UREx, we imaged whole-mount embryos following *oskar* smFISH using complementary super-resolution imaging modalities. While structured illumination microscopy (SIM) could not resolve the nanoscale organization of *oskar* RNA in the posterior condensates, STED (25% depletion laser) and lifetime-based Tau-STED (20% depletion laser) revealed clustering of RNPs within the condensates, similar to that of UREx (Fig. 4C). Note that, while the SIM and STED images were acquired with a high numerical aperture (NA) objective (N.A 1.40), the whole-mount UREx gels could only be imaged with low NA air objectives (N.A 0.75) owing to large embryo volume post-expansion.

**Fig. 4:**
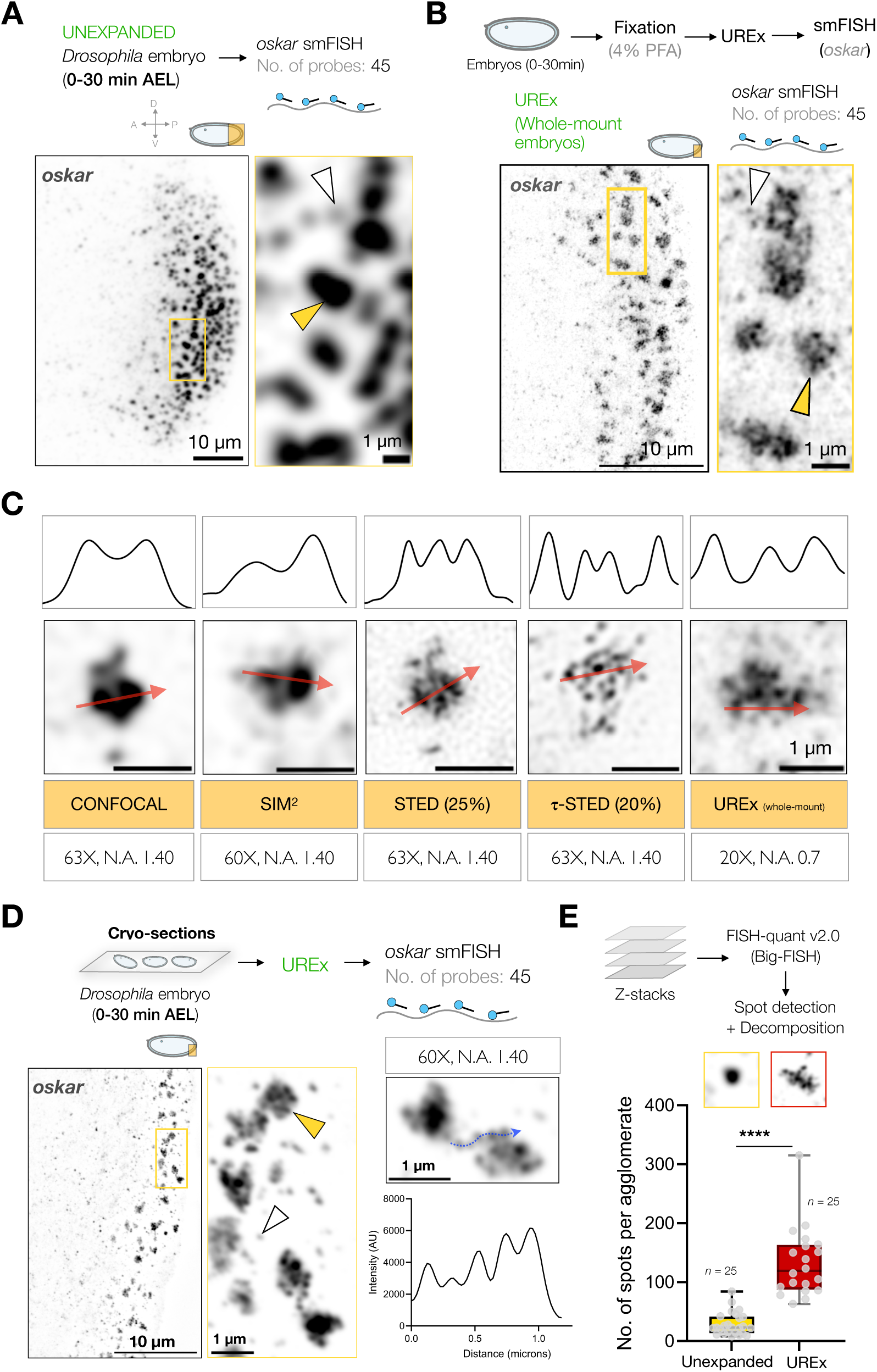
UREx resolves nanoscale organization in *oskar* RNP condensates in *Drosophila* embryos. (A) Representative confocal image of *oskar* mRNA smFISH with 45 probes (Atto 565, grey) spanning the whole mRNA at the posterior pole of early *Drosophila* embryos (0-30 minutes After Egg Laying (AEL)). The boxed area (yellow) is magnified on the right. The yellow arrowhead indicates a high copy number *oskar* RNP condensate, the white arrow indicates a low copy number *oskar* RNP granule. (B) Representative confocal image of the posterior pole of a whole-mount early *Drosophila* embryo (0-30 minutes AEL) expanded by UREx, followed by smFISH for *oskar* mRNA using the same probes (Atto 565, grey) as panel A. Image acquired using 20X air objective, NA 0.7. The boxed area (yellow) is magnified on the right. The yellow arrowhead indicates a high copy number *oskar* agglomerate super-resolved into multiple nanoclusters, and the white arrow represents a low copy number *oskar* RNP granule. (C) Representative smFISH images of single *oskar* agglomerates in unexpanded embryos (0-30 min AEL) acquired using confocal, SIM and STED microscopy. The rightmost panel represents an UREx image. smFISH of *oskar* carried out using the same 45x probe set as in panels A and B (Atto 565, grey). The STED 775 nm depletion laser power and the microscope objectives and Numerical Aperture (NA) used are indicated below. Note that the UREx image is acquired using a 20X air objective with NA 0.7. Normalized intensity profiles spanning 1 µm (real space) along the red arrows are shown on the top. (D) Representative confocal image of the posterior pole of early *Drosophila* embryo cryo-sections (0-30 minutes AEL) expanded by UREx, followed by smFISH for *oskar* mRNA using the same probes (Atto 565) as panel A. The boxed area (yellow) is magnified on the right. The yellow arrowhead represents a resolved high copy number *oskar* RNP condensate into multiple nanoclusters, and the white arrow represents a low copy number *oskar* RNP granule. Normalized intensity line profile (blue curved arrow) for multiple *oskar* RNP nanoclusters within one condensate is shown, indicating the resolution enhancement. (E) Quantification of the number of spots detected per agglomerate using the FISH-quant (v2.0) (Big-FISH) pipeline (details in Methods section). Unpaired Student’s t-test was used for comparisons. Significance level: **** p<0.0001. n = No. of condensates. In all expansion images, the scale bars are rescaled according to the expansion factor.

To facilitate imaging of expanded embryo samples with high NA objective lens, we next made 10-15 µm cryo-sections of the embryos. Although sectioning was not very robust owing to small dimensions of the fly embryo, we managed to perform UREx on some of the cryo-sections. The thin sections upon expansion were imaged using a high NA objective (NA 1.40), which resolved these large condensates into agglomerates of multiple RNPs (Fig. 4D). We used the FISH Quant v2.0 to quantify the number of spots detected per agglomerate.

Quantification from 25 agglomerates obtained from a total of 5 embryos, revealed a 5-fold increase in spot count per agglomerate for UREx compared to conventional confocal imaging, highlighting the resolution improvement achieved using UREx (Fig. 4E).

### UREx is compatible with HCR-RNA FISH

Next, we investigated if UREx could be integrated with alternative RNA in situ hybridization strategies beyond conventional smFISH. Hybridization Chain Reaction (HCR)-based RNA *in situ* hybridization methods offer higher detection sensitivity than conventional smFISH due to the hairpin-based signal amplification mechanism^36^. In contrast to conventional smFISH, where fluorescence intensity scales linearly with probe number, HCR enables the hybridization-driven assembly of fluorescent hairpins at target RNA sites, thus increasing the signal output, enabling detection of low abundance transcripts in thick tissues. Physical magnification of the sample in ExM results in cytoplasmic decrowding which facilitates the signal amplification step of HCR^36^. Moreover, the latest version HCR v3.0 uses split-initiator probes resulting in automated background suppression^37^(Extended Data Fig. 7A). We combined UREx with HCR FISH v3.0 against *oskar* RNA and were able to visualize *oskar* transcripts in the whole-mount early embryos (Extended Data Fig. 7B-C). Unlike conventional HCR protocol, using only five split-initiator probe pairs and three hours of signal amplification was sufficient to detect *oskar* signal, underlining the detection sensitivity of HCR FISH in UREx samples. Moreover, the clustering of *oskar* RNPs into agglomerates was also resolved using HCR-FISH when combined with UREx, as compared to conventional confocal imaging (Extended Data Fig. 7C).

## Discussion

Our results demonstrate that UREx can retain RNAs along with cellular ultrastructure, enabling super-resolution imaging of RNPs and RNP condensates in their native cell or tissue context. RNP granule condensates are dynamic hubs of RNA metabolism coordinating diverse processes including RNA processing, transport, storage, translation, decay, etc.^1^. Despite their functional relevance, most RNP condensates possess dimensions that are below the diffraction limit of conventional light microscopy, typically ranging from tens to a few hundred nanometers, limiting visualization of internal organization of RNAs and associated proteins. Examples include, transport granules^38^, P-bodies, transcription condensates^39^, germ granules^40^, mitochondrial RNP granules^41^, etc. Emerging evidence suggests that condensates are not compositionally homogenous and can show internal architectures such as, core–shell^20^, layered organization^42^ and nanoscale homotypic clustering^40^. Therefore, it is essential to interrogate the spatial organization within condensates and its biological significance. Although U-ExM has been applied for visualization of cytoskeletal organization^43^, organelle ultrastructure^11^, unicellular parasites^44^, viral assemblies^45^, studies to visualize RNA organization and RNP granule condensates are rare. Cirilo et al. performed U-ExM on stressed HeLa cells and observed that the core scaffold G3BP1 and stress granule (SG) component UBAP2L occupy distinct, non-overlapping patterns in SGs^46^. However, in this study immunofluorescence (IF) was used to visualize protein components of the SG condensate and RNA was not probed for. With UREx, we were able to preserve RNAs and visualize their spatial organization in nuclear paraspeckles and the cytosolic *oskar* transport granules, while preserving their native ultrastructural context.

During sample fixation, we have observed that supplementing PFA with low concentrations of GA improves RNA retention, particularly for *Neat1* (Extended Data Fig. 3). GA is a homobifunctional crosslinker that is commonly used in electron microscopy sample fixation due to its ability to promote extensive, stable cross-linking compared to that of FA. Despite its strong crosslinking capacity, GA fixation is generally less favoured for fluorescence microscopy, owing to high autofluorescence that compromises image quality^24,25^. Moreover, it has been shown that in U-ExM, GA is not optimal for structural preservation of isolated centrioles but at low concentrations with FA preserves mitochondrial ultrastructure^11^. Therefore, PFA fixation was preferred for UREx as well and GA supplementation was employed only when absolutely required.

ExM approaches have classically leveraged the highly ordered organization of protein assemblies to evaluate isotropic expansion and ultrastructure preservation. In particular, structurally conserved macromolecular complexes, such as NPCs (eightfold rotational symmetry)^23^ and centrioles (ninefold microtubule triplet-based symmetry)^11^, have served as reference standards. However, analogous RNA assemblies with structural symmetry are rarely observed in cells, making the validation of RNA ultrastructure preservation challenging. Because UREx was specifically developed to retain and resolve nanoscale RNA organization, we sought a biologically relevant RNA architecture that could serve as a benchmark for structural preservation. Consequently, we exploited the highly ordered spatial orientation of *Neat1* lncRNA in paraspeckles. The long isoform of *Neat1* is recognized as an architectural RNA that organizes the core-shell arrangement of the paraspeckles^21^. Probing the domains of *Neat1* RNA with UREx revealed the ‘looped’ arrangement of the RNA with the 5’ and 3’ extremes projecting outward and the middle domain embedded in the core of the condensates, benchmarking our technique.

Another example of an mRNA with an architectural role in condensate assembly is the maternal RNA, *oskar*, in *Drosophila melanogaster*^14^. We have shown that *oskar* RNA acts as an architectural scaffold and together with Bruno nucleates *oskar* granule assembly in the oocyte^38^. With UREx, we discover that multiple *oskar* granules cluster into ‘agglomerates’ in the germ plasm of fertilized eggs (Fig. 4). The molecular mechanism and the functional importance of this clustering in germline development remains to be investigated. Moreover, the organization of key RBPs such as, Staufen, Bruno, PTB, etc within these agglomerates can be further probed leveraging the robust detection sensitivity and resolving power of UREx. We have demonstrated that UREx is compatible with at least two hybridization-based RNA detection methods, smFISH and HCR-FISH. While smFISH ensures high fidelity RNA detection, HCR-FISH can be applied to detect low abundant transcripts in thick tissue samples with high sensitivity. Furthermore, we performed UREx on whole-mount embryos, suggesting its applicability to image tissues or intact organisms that are dimensionally not amenable to sectioning.

The available ExM approaches that are shown to retain RNAs include, Click-ExM and Magnify. Click-ExM relies on biorthogonal Click chemistry to metabolically or chemically label individual classes of biomolecules to the gel matrix^9^. Magnify, on the other hand, employs Methacrolein-based hydrogel formation to universally retain all biomolecules^10^. Despite claiming RNA retention, neither approach provides examples of super-resolved RNA-containing cellular structures nor benchmarks RNA ultrastructure conservation using endogenous RNP assemblies. In contrast, we demonstrate that UREx preserves RNA-protein ultrastructure in standard chemically-fixed samples and U-ExM-compatible hydrogel chemistry, without the requirement of biomolecule-specific clickable reagents, metabolic labelling strategies or proprietary anchoring chemistry which may not be readily accessible to standard laboratories. Moreover, instead of using commercially labelled probe sets, we labelled our smFISH oligonucleotide probes in the laboratory at the 3’-end with Atto dyes using Terminal Deoxynucleotidyl Transferase (TdT)-based ddUTP addition^47^, which significantly reduces the cost. Thus, the absence of specialized RNA-anchoring chemistry^6, 9^ or expensive chemically modified probes^7^, together with the compatibility with in-house labelled probes, further underscores the ease of adapting the UREx workflow and its overall cost-effectiveness.

In future, UREx can be potentially combined with multiplexed RNA FISH to study spatial analysis of gene expression at unprecedented resolution. Furthermore, combining UREx samples with optical super-resolution imaging modalities such as STED, SIM can further enhance the attainable resolution.

## Extended Data Figures

**Extended Data Fig. 1:**
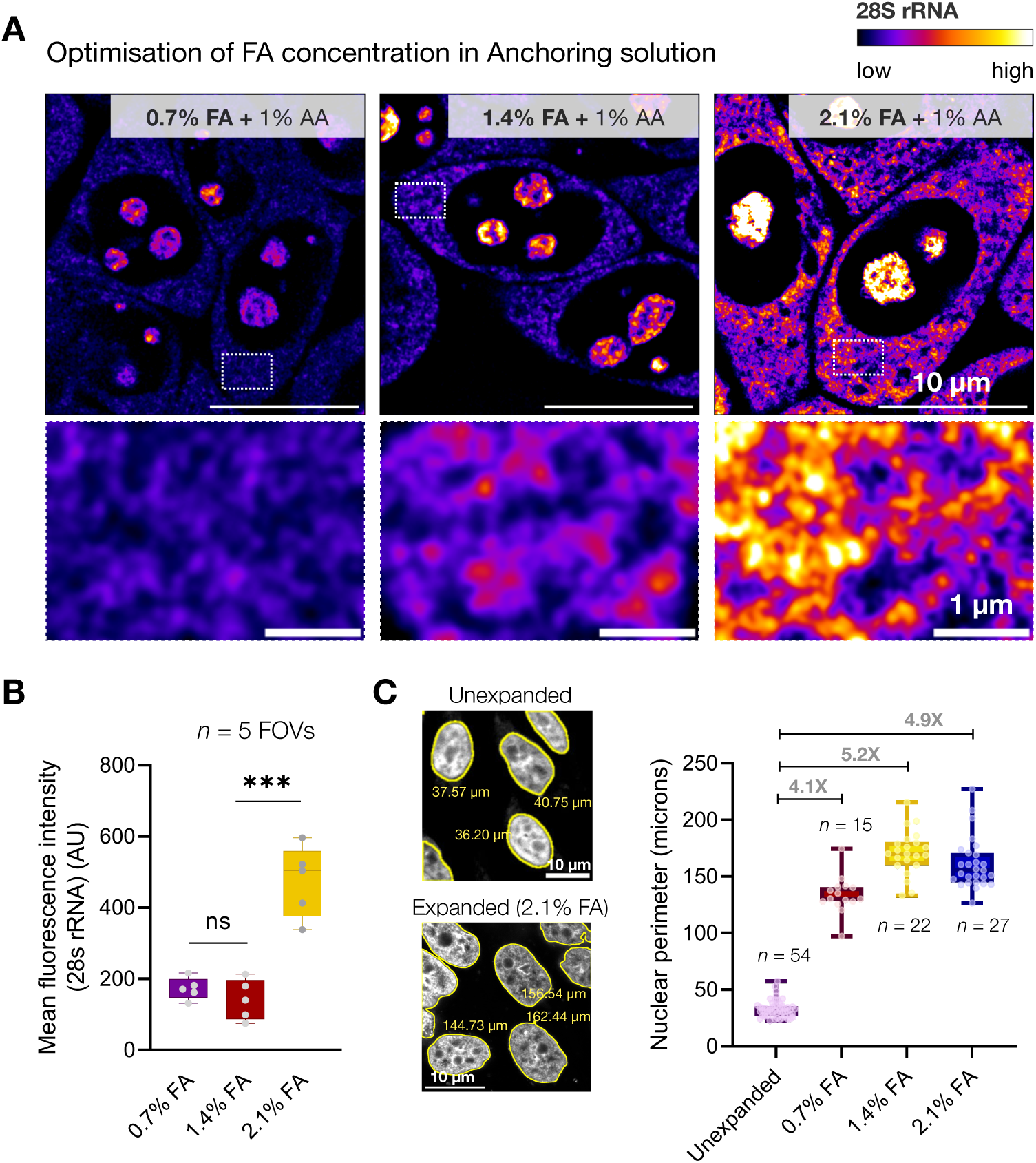
Optimization of the UREx protocol. (A) Representative confocal images of smFISH with 28S rRNA (Atto 633) probes in expanded Chinese Hamster Ovary (CHO) cells anchored with 1% AA and 0.7%, 1.4%, or 2.1% FA. (B) Comparison of mean fluorescence intensity of 28S rRNA signal in expanded CHO cells under different anchoring conditions. Unpaired Student’s t-test was used for comparisons. n = number of field of views (FOVs). *** p<0.001, ns (non-significant). (C) Comparison of the nuclear perimeter in unexpanded and expanded CHO cells under different anchoring conditions. n = number of nuclei. The expansion factors are indicated above the box plot. Representative Hoechst-stained nuclei and perimeter (yellow) of unexpanded and (2.1% FA + 1%AA) anchored cells are shown on the left. Scale bars are adjusted according to the expansion factor.

**Extended Data Fig. 2:**
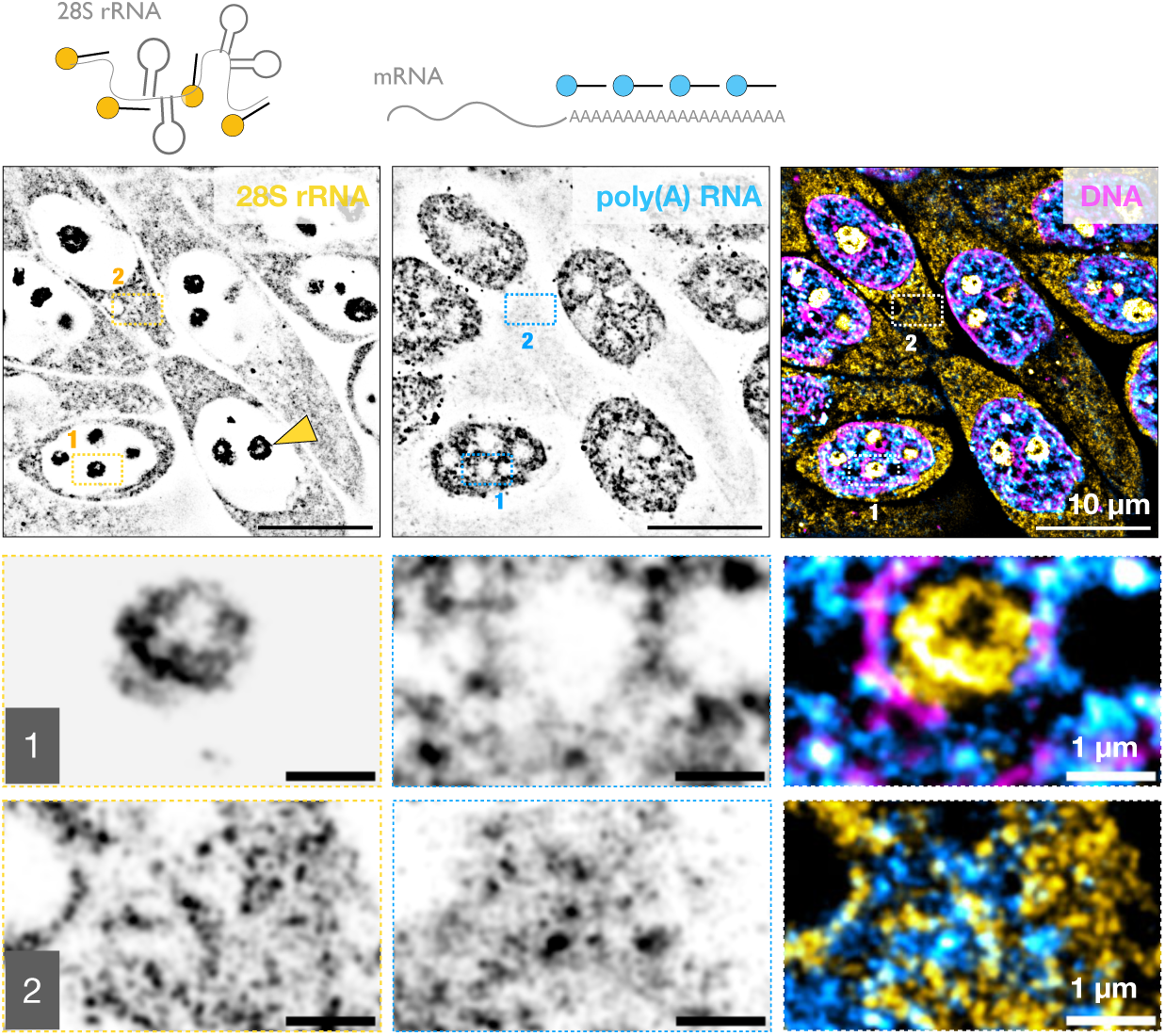
Visualizing the cellular pool of rRNA and poly(A) RNA using UREx. Representative confocal images of CHO cells expanded by UREx with 28S rRNA (yellow), poly (A) RNA (cyan) visualized by smFISH with oligo-dT (Atto 565) and 28S rRNA (Atto 633) antisense probes, respectively; DNA (magenta) stained using Hoechst. Boxed areas 1 (nucleoplasm) and 2 (cytoplasm) are enlarged below. Scale bars are adjusted according to the expansion factor.

**Extended Data Fig. 3:**
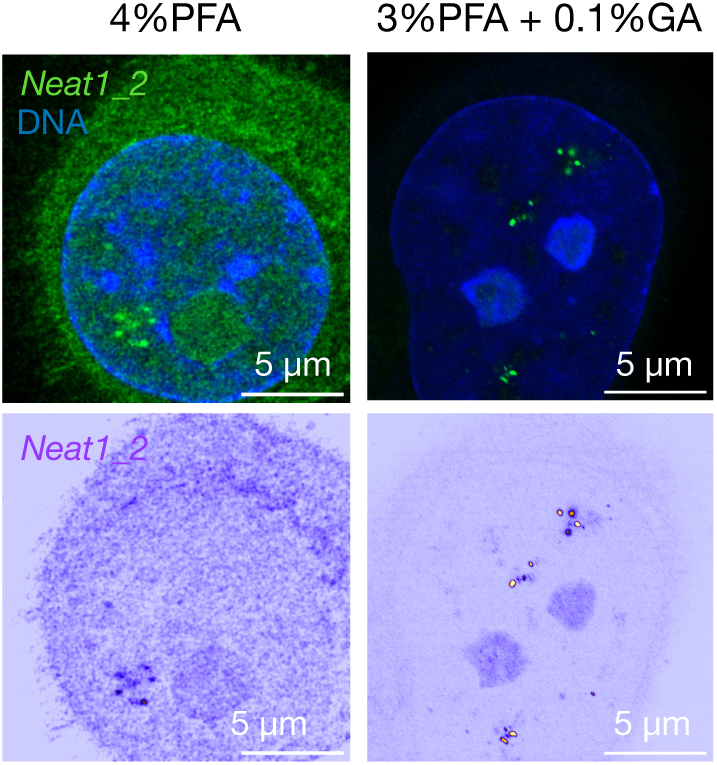
Optimization of the fixation procedure to retain paraspeckles in the gel. Confocal images of *Neat1_2* smFISH using 5’+Mid probes (60X) (Atto 633, green) in UREx expanded HeLa cells fixed with 4% PFA (left) or 3% PFA+0.1% GA (right). Lower panels are heat-map representation of the *Neat1* smFISH signal to depict the improvement in signal-to-noise ratio (SNR) upon GA spiking. DNA stained using Hoechst. Note that the PFA+GA fixed cell is the same cell as shown in Figure 1B. Scale bars are adjusted according to the expansion factor.

**Extended Data Fig. 4:**
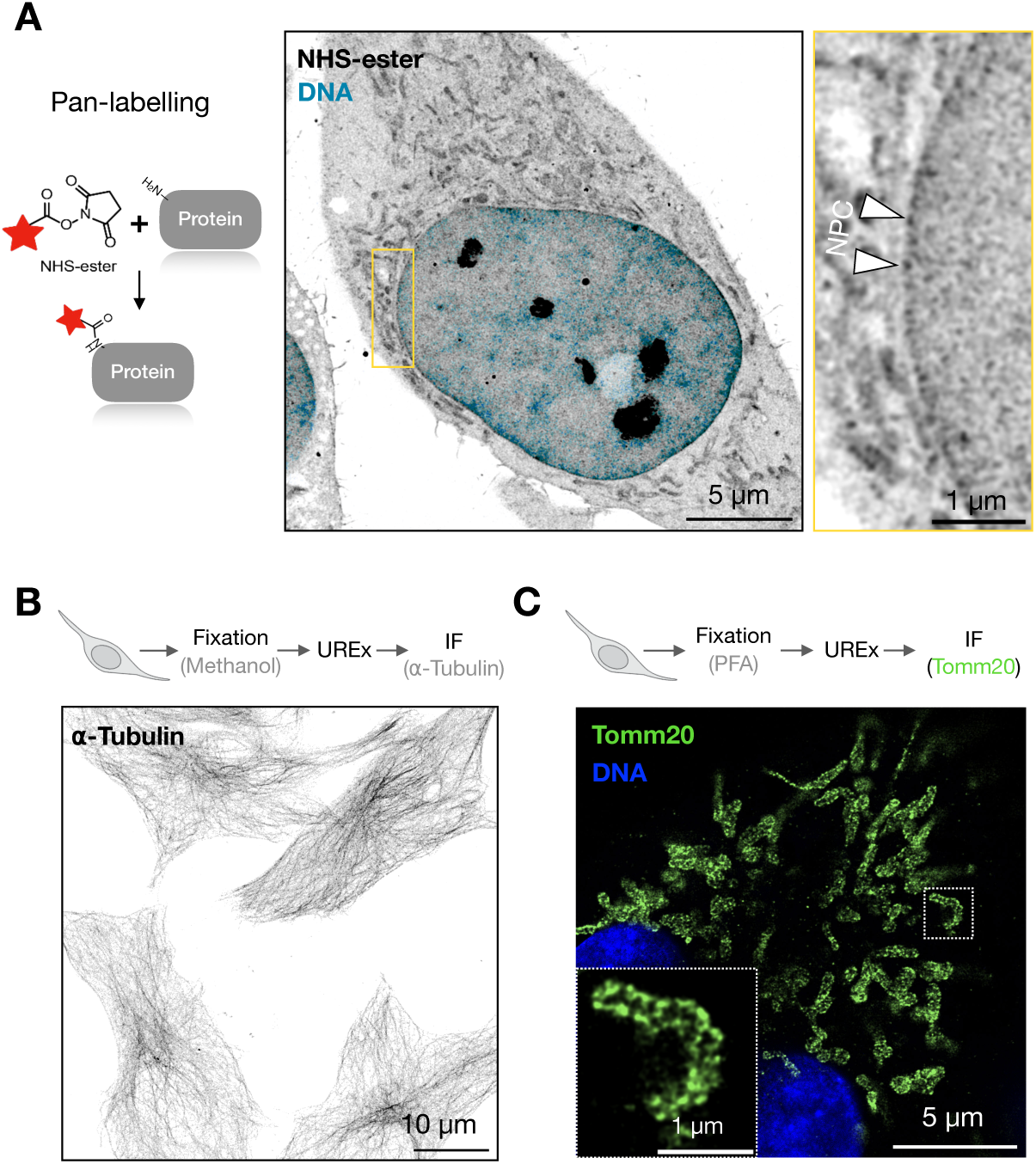
UREx protocol preserves native cellular ultrastructure and specific protein epitopes. (A) Atto 488 NHS-ester and Hoechst staining of expanded HeLa cells anchored using 2.1% FA + 1% AA. Boxed area (yellow) is magnified on the top right; arrows indicate Nuclear Pore complexes (NPC). (B) Visualization of the microtubule cytoskeleton in methanol-fixed Hela cells by IF using anti-Tubulin FITC antibody (1:250). DNA is visualized by Hoechst staining. (C) UREx coupled with IF against outer mitochondrial membrane protein Tomm20 (1:1000) in PFA-fixed HeLa cells and visualized using Alexa Fluor 488 conjugated secondary antibody (1:1000). DNA is visualized by Hoechst staining. Scale bars are adjusted according to the expansion factor.

**Extended Data Fig. 5:**
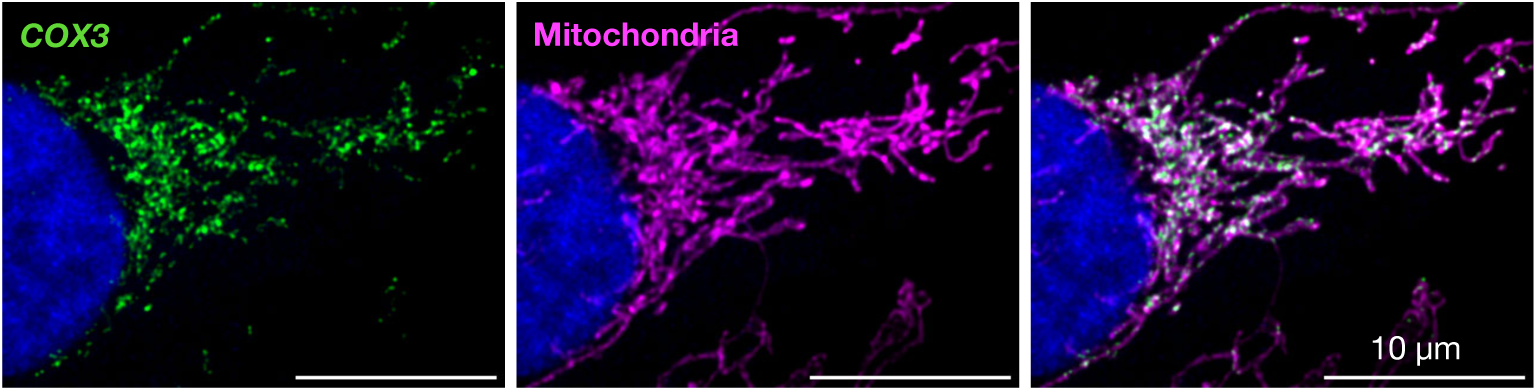
Distribution of mitochondrial transcripts in unexpanded Hela cells. Representative confocal images of *COX3* smFISH (Atto 565, green) in unexpanded Hela cells. Mitochondria are labelled using Mitotracker (magenta) and DNA (blue) is stained by Hoechst.

**Extended Data Fig. 6:**
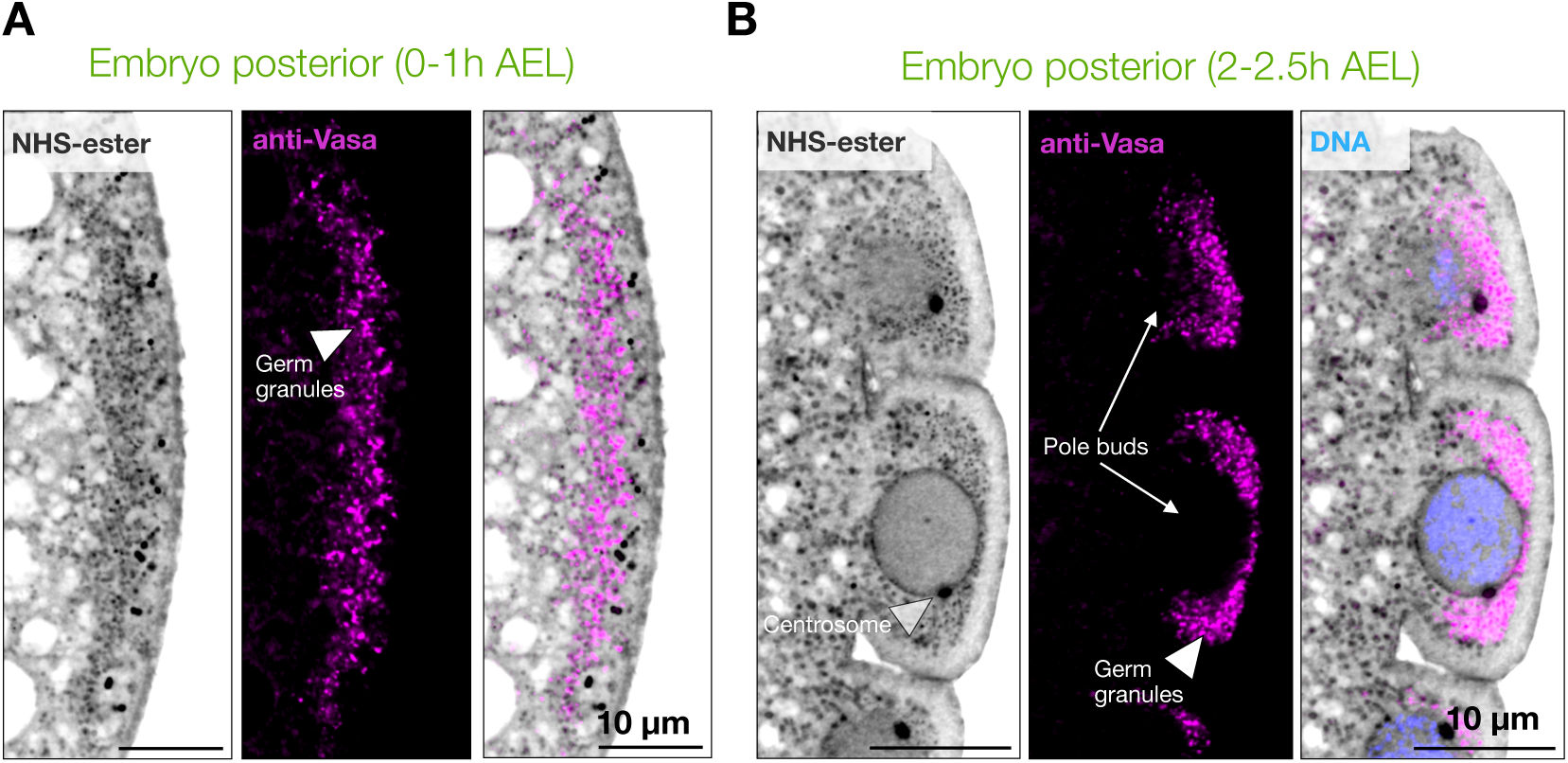
Visualization of germ granule condensates in *Drosophila* embryos using UREx. (A-B) Representative confocal images of whole-mount *Drosophila* embryos 0-1 h AEL (A) and 2-2.5 h AEL (B) after UREx, stained with anti-Vasa primary antibody (1:200) and Alexa Fluor 488 conjugated secondary antibody (1:200). Pan-labelling was performed with Atto565 conjugated NHS-ester, and DNA visualised with Hoechst. Scale bars are adjusted according to the expansion factor.

**Extended Data Fig. 7:**
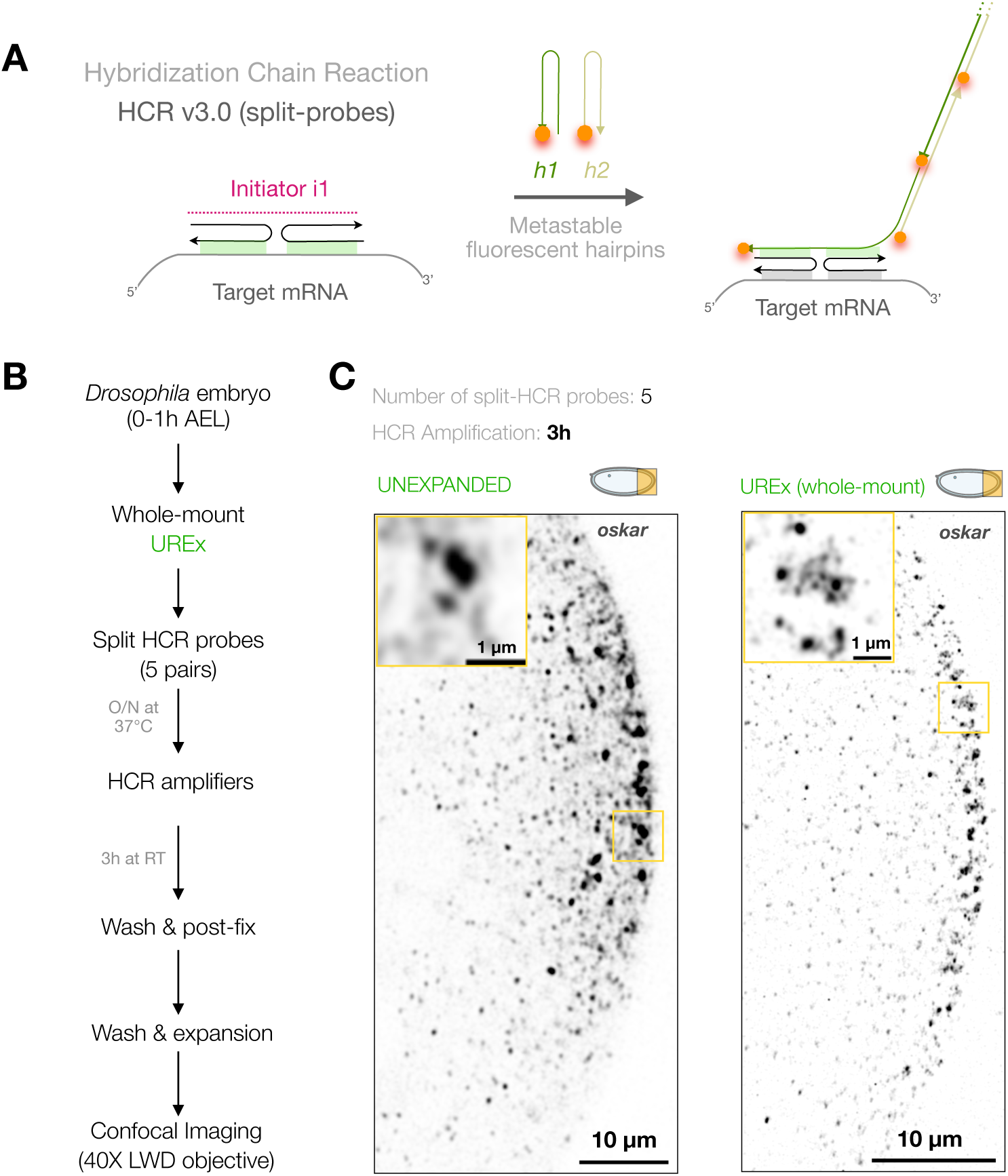
UREx coupled with HCR-RNA FISH resolves *oskar* RNP condensates into nanoclusters in young embryos. (A) Schematic representation of the principle of HCR-FISH v3.0 with split-initiator probes. (B) Schematic workflow of UREx coupled with HCR-FISH v3.0 on whole-mount early embryos (0-1h AEL). (C) Representative confocal images of whole-mount *Drosophila* early embryo posterior pole unexpanded (left) and expanded by UREx (right), followed by HCR-FISH v3.0 for *oskar* mRNA with five split-initiator probe pairs. Alexa Fluor 546-labelled B2 hairpin amplifiers were used for detection, and signal amplification was carried out for 3 h. Boxed regions (yellow), magnified on top left, show individual condensates resolved into nanoclusters. The data show that only 5 probe pairs and 3 h of signal amplification are sufficient for detection and resolution of condensates into nanoclusters by UREx. Scale bars are adjusted according to the expansion factor.

## Supporting information

List of all chemical reagents and antibodies used

List of smFISH probes used in the study

Sequences of HCR split-initiator probes used in the study

## Acknowledgements

We acknowledge the Department of Bioscience and Biotechnology, IIT Kharagpur and DST-FIST for the FV3000 confocal microscopy facility. We acknowledge Budhaditya Mukherjee, School of Medical Science and Technology, IIT Kharagpur for access to the Leica Stellaris microscope. We acknowledge the central instrumentation facility of IIT Kharagpur and Arindam Mondal for access to the STED and SIM super-resolution microscopes. We thank Budhaditya Mukherjee, Shubhangi and Sourav Dutta for assistance with embryo cryo-sections at the Leica cryostat. We are grateful to Anne Ephrussi, Julia Mahamid, Florence Besse and Jeffrey Chao for their critical comments on the manuscript. We thank Piyali Mukherjee for the anti-Tomm20 antibody. We sincerely thank Anne Ephrussi for anti-Vasa, anti-Tubulin antibodies and *oskar* smFISH probe sets. M.B acknowledges the funding from IIT Kharagpur and ANRF-India.

## Author Contributions

P.H.D and M.B conceived the study. J.M, P.H.D and M.B designed the experiments, interpreted the results and wrote the manuscript. J.M and P.H.D performed the experiments and analyzed the data. S.B and P.P adapted the FISH-quant pipeline for smFISH spot detection.

## Materials and Methods

### Cell culture and fixation

HeLa and CHO cells were cultured in DMEM supplemented with 10% FBS, 1% Penicillin-Streptomycin, 1% Glutamax. Cells were maintained in T-25 flasks and seeded on 12 mm cover slips placed in 24 well plates at a density of approximately 1.5 x 10^5^ cells per well for experiments. Cells were fixed using 4% paraformaldehyde, unless mentioned otherwise. To visualize mitochondria, Mitotracker DeepRed (Thermo Scientific) was used at a working concentration of 100 nM. When the seeded cells reached 70% confluency, the old media was removed and fresh media containing the mitotracker dye was added and the cells were incubated in the 37°C, 5% CO_2_ incubator for 20-30 minutes. Following this, 3% paraformaldehyde (PFA) + 0.1% Glutaraldehyde in PBS was used to fix the cells for 15 min at room temperature (RT) as previously described^8^ and processed for smFISH/UREx. For *Neat1_2* UREx-smFISH, cells were also fixed in 3% paraformaldehyde (PFA) + 0.1% Glutaraldehyde in PBS.

### Fly maintenance

*Drosophila melanogaster* stocks were maintained at 25°C on standard cornmeal agar. *w*^1118^ wild-type flies were used for all experiments. For egg laying, 2–5-day old female flies were transferred to food vials with double as many male flies with fresh yeast for 2 days and subsequently transferred to cages for egg laying and embryo collections.

### *Drosophila* Embryo collection

2–5-day old flies were fed with yeast powder for at least 2 days and transferred to cages attached with 35mm apple agar plates with a drop of yeast paste. After one or two rounds of pre-laying (30 min), eggs were harvested from plates after 0–30 min of egg laying at 25°C. The apple agar plates were detached from the cage and 50% bleach solution was applied for 2 minutes for dechorionation. Meanwhile, the fixative solution (1:1 solution of 4% PFA and heptane) was prepared in a 1.5 ml microcentrifuge tube. The embryos from the plates were transferred to a sieve and extensively washed with dH_2_O to remove the excess bleach. Subsequently the embryos were transferred using a thin brush to the freshly prepared fixative solution. Fixation was performed at RT for 20-25 minutes with 1000 rpm shaking. After fixation, the lower layer in the tube containing PFA was gently removed by using a Pasteur pipette and an equal amount of 100% methanol was added and vortexed at the highest speed for 1-2 minutes to remove the vitelline membrane. The devitellinized embryos, settled at the bottom of the tube, were collected and rinsed with 100% methanol for 2-3 times. The embryos were then finally stored in 100% methanol in -20°C for later use.

### smFISH probe labelling

#### Dye conjugation to NH_2_-ddUTP

Atto 633, Abberior STAR 580, Atto 565 and Atto 488-NHS esters were reconstituted in anhydrous DMSO at a final concentration of 20 mM. NHS esters were conjugated to amino-11-ddUTP by using a 2-fold molar excess of dye-NHS-ester in the presence of 0.1 M NaHCO3 pH 8.3 for 2 hours at RT in dark. The reaction was stopped with 1 M Tris HCl pH 7.4 and the final concentration was adjusted to 5 mM by using nuclease-free water.

#### Labelling of oligonucleotide probe mix

Probe sequences used for smFISH experiments are described in the Supplementary Table 1. 18-22 nucleotide long non-overlapping DNA oligos (Barcode Bioscience) was selected using the smFISHprobe_finder.R script and reconstituted to 250 μM in nuclease-free water. Using the Terminal Deoxynucleotidyl Transferase (Thermo scientific) enzyme, the probes were enzymatically conjugated to the Atto 633, Atto 565 or Atto 488-ddUTP by overnight incubation at 37°C. The labelled oligos were recovered by precipitation using absolute ethanol, sodium acetate pH 5.5 and linear acrylamide and finally dissolved in nuclease-free water. The concentration of the labelled probe mix and the degree of labelling were measured in a Nanodrop spectrophotometer as described previously^47^.

### smFISH in unexpanded samples

#### smFISH in cultured cells

Cells were cultured and fixed as described above. Subsequently, cells were permeabilized with 70% ethanol overnight at 4°C. Next day, the cells were washed twice with 0.1% PBT for 10 minutes at RT and prehybridization solution (10% formamide, 2X sodium citrate in dH_2_O) was added and incubated for at least 5 minutes at RT. Meanwhile, a humid chamber was made using a petri plate and 30 μl of hybridization solution (10% dextran sulphate, 10% formamide, 2X sodium citrate in dH_2_O) with 2.5 nM/probe was added on a parafilm strip. The cover slip was transferred from the 24 well plate to the parafilm, upside down, using sharp tweezers. Incubation was carried out at 37°C for 3-4 hours in the dark. Next the cells were washed twice with the prehybridization solution on the parafilm itself for 15 minutes each at 37°C in the dark. Subsequently, the cover slip was transferred back to the 24 well plate and the cells were washed with 0.1% PBT for 10 minutes at RT on a shaking rocker. The nuclei were stained with 1 drop of NucBlue (Thermo scientific) in 1 ml 0.1% PBT for 15 minutes and cells were subsequently washed for 10 minutes with 0.1% PBT on the shaking rocker at RT. Cover slips were mounted on a fresh glass slide using Immu-Mount (Fisher scientific) and imaged.

#### smFISH in whole mount embryos

Embryos were collected, fixed and preserved as described above. After removal of methanol, the stored embryos were first rinsed and then rehydrated by washing with 0.1% PBT thrice for 15 minutes each on the shaking rocker at RT. Next prehybridization solution (2X sodium citrate, 10% formamide, 0.1% Tween-20 in dH_2_O) was added and incubated at 37°C for 30 minutes with shaking (1000 rpm) in a thermomixer. Next the prehybridization solution was removed and 200 μl of hybridization solution (2X sodium citrate, 10% formamide, 0.1% Tween-20, 2mM vanadyl ribonucleoside, 100 μg/ml sheared salmon sperm DNA, 10% dextran sulphate, 20 μg/ml BSA in dH_2_O) containing 2.5 nM/probe was added and incubated overnight at 37°C with 1000 rpm shaking in a thermomixer. The following day, the hybridization solution was removed and the embryos were washed with the prehybridization solution twice for 15 minutes each at 37°C with 1000 rpm shaking. Subsequently, a 0.1% PBT wash was done at RT on the shaking rocker. DAPI staining was done in 0.1% PBT for 15 minutes at RT and embryos were washed twice with 0.1% PBT for 10 minutes each at RT. Next, mounting media (80% Glycerol, 2% Propyl Gallate) was added and the embryos were incubated at 4°C for at least 2 hrs. Subsequently, the embryos were mounted and arranged on a fresh slide and imaged.

### Cryo-sectioning of *Drosophila* embryos

*Drosophila* embryos were harvested and fixed as mentioned above. Next, they were incubated overnight in a cryo-protection solution (3.7% (v/v) formaldehyde, 30% sucrose in 1X PBS) at 4°C. Then, the embryos were stained with bromophenol blue solution (for visualisation in the frozen block) and arranged under a stereo microscope in a cryo-mould whose surface was coated earlier with a double-sided tape. The excess solution was then removed followed by gentle addition of Tissue Freezing Medium (TFM, SIGMA) on the tissue. The mould was kept inside a -80°C freezer for at least 1 hour for freezing. 10-15 µm sections were cut on a Leica cryostat and transferred to the surface of poly D-Lysine (PDL)-coated 12 mm coverslips. The tissue sections were post-fixed with 4% PFA for 20 minutes at RT and washed with 0.1% PBT for 10 minutes and immediately processed for the anchoring step of UREx or stored in -20 °C freezer for 3-4 days.

### UREx in cultured cells and cryo-sections

Following fixation, cover slips with cells or cryo-sections were incubated with UREx anchoring solution (2.1 % FA + 1% AA) in 1x PBS at 4°C overnight. The anchoring solution was supplemented with 0.5 mM vanadyl ribonucleoside RNase inhibitor to protect the cellular RNAs from degradation. Gelation was performed by incubating the coverslips with cells/cryo-sections facing down in 35 μl of monomer solution (19% wt/wt Sodium Acrylate (SA), 10% (wt/wt) Acrylamide (AA), 0.1% (wt/wt) Bis-Acrylamide (BA) in 1x PBS supplemented with 0.5% APS and 0.5% TEMED) on a parafilm-coated slide in a humid chamber at 4°C for 5 minutes, and then at 37°C for 1 hour in the dark. Following gelation, denaturation was performed by removing the coverslip containing gel from the parafilm using sharp tweezers and incubating with denaturation solution (200 mM SDS, 200 mM NaCl, and 50 mM Tris-Cl (pH 9) in dH2O) for 15 minutes on a shaker in a 35mm petri dish at RT (to detach the gel from the coverslip). The gel was then incubated at 95°C for 1 hour in a 1.5 ml microcentrifuge tube with 1 ml of fresh denaturation solution. All UREx buffers were prepared in nuclease-free ultrapure water. To visualize mitochondrial RNAs in HeLa cells, the denaturation step was performed at 70°C for 1 hour ^8^. After removing the denaturation solution, the gel was kept in a 90 mm petri dish containing H_2_O enough to cover the gel for 2 h to overnight at RT for removal of SDS and the first round of expansion. H_2_O was changed twice in a 30 min interval. After that, the gel was shrunk in 1x PBS and gel punches obtained using the base of a 1000 μl pipette tip and proceeded for smFISH/HCR-FISH/IF/NHS-pan-labelling. Final expansion was carried out in nuclease-free ultrapure water.

### UREx in whole-mount *Drosophila* embryos

*Drosophila* embryos were harvested and fixed as described above. In case the sample was stored in dehydrating conditions at -20°C, rehydration was performed using 0.1% PBT thrice for 15 minutes each on the rocking shaker at RT. Following this, the sample was incubated in UREx anchoring solution at 4°C overnight. Under a stereo microscope, the embryos were manually arranged on a 12 mm coverslip and proceeded for the gelation step as described above. Denaturation was performed at 95°C for 1 hour.

### UREx-smFISH

smFISH was carried out by incubating the gel punches in 1 ml pre-hybridization solution (2X sodium citrate, 10% formamide, 0.1% Tween-20 in dH_2_O) at 37°C for 30 minutes with rotation (1000 rpm) in a thermomixer. The prehybridization solution was removed and 200 μl hybridization solution (2X sodium citrate, 10% formamide, 0.1% Tween-20, 2 mM vanadyl ribonucleoside, 100 μg/ml sheared salmon sperm DNA, 10% dextran sulphate, 20 μg/ml BSA in dH_2_O) containing 25 nM/probe and incubation was done at 37°C overnight with shaking (1000 rpm) in thermomixer. After the removal of the hybridization solution, the gels were washed twice with prehybridization solution at 37°C for 15 minutes each with rotation (1000 rpm) and 0.1% PBT at RT with shaking. Next, the gels were stained with Hoechst (200 μg/ml) in 0.1% PBT for 15 minutes, followed by washing with 0.1% PBT at RT with shaking.

For NHS-ester pan-labelling, Atto488-NHS ester solution was prepared in 0.1% PBT at a concentration of 5 μM and gels incubated for 30 minutes at RT with shaking. Lastly, the gel punches were kept in a 90 mm petri dish containing dH2O for 1-3 hours for final expansion before imaging.

### UREx-smFISH + IF

UREx-smFISH was carried out as described above. Following this the gel punch was post fixed using 4% paraformaldehyde at RT for 20 minutes with shaking (1000 rpm) in a thermomixer. The sample was then washed 3 times for 5 minutes each in 0.1% PBT at RT on a shaking rocker. Subsequently, the gel punch was incubated with primary antibody (Tomm20, 1:1000) diluted in blocking buffer (1% BSA in 0.1% PBT) for 1 hr at RT on the shaking rocker. After incubation, the sample was washed thrice for 10 minutes each with 0.1% PBT at RT on the shaking rocker. Next, the sample was incubated with fluorophore conjugated secondary antibody (1:1000) diluted in the blocking buffer (1% BSA in 0.1% PBT) for 2 hrs at RT on the shaking rocker followed by 3 washes using 0.1% PBT at RT. samples were then stained with DAPI and transferred to dH_2_O for 1-3 hours for final expansion before imaging. For IF alone, all the steps were identical except the incubations with primary antibody (Tomm20, 1:1000; anti-Tubulin FITC, 1:250) were carried out overnight at 4°C.

### UREx-HCR-FISH v3.0 in whole-mount embryos

For HCR-RNA FISH, UREx gel punches were pre-hybridized with 1 ml pre-heated (37°C) HCR probe-hybridization buffer (probe HB) (30% formamide, 5× sodium citrate (SSC), 9 mM citric acid (pH 6.0), 0.1% Tween 20, 50 μg/mL heparin, 1× Denhardt’s solution, 10% dextran sulfate) for 30 minutes at 37°C with rotation (1000 rpm) in a thermomixer. The probe solution was prepared by mixing 0.4 pmol of each split-probe set (1 μl from 0.4 μM stock diluted in probe HB) in 200 μl of probe HB. Hybridization with the probe solution was carried out at 37°C overnight with shaking (1000 rpm). Post-hybridization, the gel was washed with pre-heated (37°C) probe wash buffer (probe WB) (30% formamide, 5× sodium citrate (SSC), 9 mM citric acid (pH 6.0), 0.1% Tween 20, 50 μg/mL heparin). Gels were washed 4x 15min with 0.5 ml of probe WB at 37°C with rotation (1000 rpm) and washed 2x 5min with 1ml 5x SSCT buffer (5× sodium chloride sodium citrate (SSC), 0.1% Tween 20) at RT with shaking.

Meanwhile, the amplification buffer (AB) (5× sodium citrate (SSC), 0.1% Tween 20, 10% dextran sulfate) was equilibrated at RT, and the gels were pre-amplified with AB for 10 minutes at RT with shaking. Separately, 15 pmol of hairpin h1 and 15 pmol of hairpin h2 were prepared by snap cooling 5 μL of 3 μM stock (heat at 95°C for 90 seconds and cool to room temperature in a dark drawer for 30 min). Next, the hairpin solution was prepared by mixing the hairpins in 200 μl AB and added to the gels. The gel was then incubated overnight at RT with shaking in the dark. The gel was subsequently washed with 5x SSCT for 2 x 5 min, 2 x 30 min, 1 x 5 min at RT with shaking in the dark, and post-fixed with 4% PFA for 20 minutes followed by final washing twice with 5x SSCT at RT for 10 minutes each. After that, the gel was kept in a petri dish containing dH_2_O for final expansion for 1-3 hours at RT in the dark.

### Gel Mounting and image acquisition

For confocal imaging the gels were mounted on PDL-coated glass-bottom dishes (20 mm wide glass surface), with the tissue/cell surface facing the glass surface.

Unexpanded samples of cultured cells/embryos/ovaries as well as UREx gels with cells/tissue sections were imaged using a 63X oil immersion objective lens (NA 1.4) or 60X oil immersion objective lens (NA 1.4) in Leica STELLARIS or Olympus FV3000 laser scanning confocal microscope. High resolution images were acquired with Nyquist sampling and deconvolved using the lightning module of Leica STELLARIS or Fiji plugins Diffraction PSF 3D (to calculate theoretical point spread function (PSF) and Iterative Deconvolve 3D for 5-10 iterations. UREX-smFISH whole mount embryos were imaged using 20X air objective (NA 0.75) in Leica STELLARIS microscope. For imaging of whole mount embryos after IF or UREx-HCR FISH, a 40X long working distance objective (NA 0.7) was used and images acquired in the Olympus FV3000 laser scanning confocal microscope under aforementioned settings. Note that for expansion microscopy, owing to volumetric dilution of the fluorophores, higher laser power was used and higher pixel dwell time was required for image acquisition to capture sufficient photons.

### STED and SIM imaging

Paraspeckles in HeLa cells (*Neat1_2* smFISH probe combinations labelled with Atto 633 or Abberior STAR 580) were imaged using the Leica STELLARIS STED platform using a 63X oil immersion objective (NA 1.40) in the confocal mode as well as Tau-STED mode (20% intensity, 775 nm depletion laser) at a pixel size of 20 nm. For *oskar* smFISH (Atto 633) in unexpanded embryos, images were acquired in intensity mode with STED 775 nm depletion power 25% as well as Tau STED mode with 775 nm depletion laser power of 20%.

For SIM imaging of unexpanded embryos after *oskar* smFISH (Atto 633), the posterior half of the embryo was imaged using 60X oil objective (NA 1.4) of Zeiss Lattice SIM 5 Super resolution microscope with 50 ms exposure and 42 µm grating at a voxel size of 34 nm x 34 nm x 140 nm. This was followed by 3D processing of the images using the in-built SIM^2^ pipeline for 3 iterations and standard sharpness conditions.

### Image analysis

For all image representations in the figures, maximum intensity projections of z-slices, intensity line profiles, Fiji was used.

#### Quantification of 28s rRNA retention in different anchoring conditions in cultured cells

For the estimation of 28S rRNA retention in CHO cells, smFISH with 28S rRNA specific 20x antisense DNA probe was performed on the expanded cells. The 28S rRNA signal was segmented based on intensity using trainable Weka segmentation in Fiji. The measured mean fluorescence intensity was compared between the different anchoring conditions used for expansion.

#### Intensity Line profiles

To enable comparison, normalized line intensity profiles were generated for images across all modalities.

#### Quantification of expansion factor in cultured cells

For the estimation of expansion factor in cells, unexpanded and expanded cells were stained with Hoechst. For nuclear segmentation and expansion factor calculation, intensity-based thresholding of the nuclei of CHO cells was performed in Fiji using the default settings and nuclear perimeter quantified using the Fiji particle analysis tool. The measured nuclear perimeter was compared between unexpanded (n=54 nuclei) and expanded with anchoring containing 0.7% FA (n=15 nuclei), 1.4% FA (n=22 nuclei), 2.1% FA (UREx) (n=27 nuclei).

#### Quantification of nearest-neighbour distance between Neat1 domains

The three colour UREx images were quantified using a custom automated Python pipeline. The fluorescent signals for 5′, 3′, and mid regions were segmented using the multi-Otsu algorithm followed by filtering of objects less than 4 pixels, as explained earlier. To account for the discontinuity between the 5′ and 3′ shells of *Neat1*, a local maxima peak finding algorithm was applied with a minimum peak distance of 3 pixels to determine the centroids of the 5′ and 3′ shells. The Euclidean distances from 5′ to the core (mid) centroid, from 3′ to the core (mid) centroid, and from 5′ to 3′ were quantified using nearest-neighbour proximity across the entire field of view.

#### Calculation of 5′ to core (mid) edge distance in Neat1 paraspeckles

The spatial distributions of 5′-to-core (mid) distances of *Neat1* RNA were quantified using a custom automated Python pipeline. True paraspeckles’ 5′ and mid signals from maximum-intensity projections were segmented using a three-class multi-Otsu thresholding algorithm. The objects smaller than 4 pixels were filtered out. To rule out structural anisotropy, the core (mid) shell (5′) distances were measured by an edge-to-centroid nearest neighbour approach. The outer boundary of the core was calculated mathematically by taking a 1-pixel thick outline from its outer perimeter. To account for the discontinuity in the 5′ shell of *Neat1*, a local maxima peak finding algorithm was applied with a minimum peak distance of 3 pixels, to find the centroids of the 5’ shell. The Euclidean distance from the 5′ centroid to the nearest pixel of the outer boundary of the mid core was calculated.

#### Quantification of number of RNA spots per unit mitochondrial area

The spatial distribution of MTCO3 spots in mitochondria was quantified using a custom automated Python pipeline. The mitochondria were segmented using a multi-Otsu thresholding algorithm, lowest foreground thresholding was applied to account for the high intensity core and low intensity mitochondria edges. The objects less than 49 pixels were filtered out, and internal holes were filled. The total mitochondrial area was calculated and rescaled back using the expansion factor for expanded samples. MTCO3 spots were detected using the Big-FISH spot detection algorithm with a spot radius of 200 nm. The spot coordinates that colocalized within segmented mitochondrial boundaries were considered to quantify MTCO3 spots per micron^2^ in expanded and unexpanded samples. The detected MTCO3 spots and mitochondrial segmentation were visualized using Napari.

#### Spot detection using FISH-quant (v2.0) (Big-FISH) pipeline

We detected the RNA spots and quantified them using Big-FISH (FISH-quant v2.0) Python package^4849^. For quantification of the number of spots in a single agglomerate in expanded and unexpanded samples, 3D ROIs containing single agglomerates were cropped. The 3D stacks were pre-processed with standard Big-FISH (FISH-quant v2.0) pipeline and the spots were detected using adaptive thresholding with an estimated radius of 10 nm more than the voxel size. Dense spots were decomposed (α = 0.7, β = 1, γ = 5) followed by subpixel localization by gaussian fitting.

#### Estimation of Pearson’s Correlation Coefficient (PCC-Costes)

For quantifying the degree of colocalization of the *COX3* and *ND5* mRNAs using PCC, in expanded and unexpanded samples (Fig. 1D), n=12 unexpanded and n=8 expanded FOVs were analyzed using the BIOP-JACoP fiji plugin. For quantifying colocalization of the 5’ and 3’ ends of *oskar* mRNA in expanded and unexpanded samples (Fig. 2E), n=5 unexpanded and n=6 expanded images were analyzed using BIOP-JACoP Fiji plugin. Otsu algorithm was applied for thresholding. Significance of colocalization was estimated using Costes randomization (100 shuffles over 5 pixel-sized blocks).

## Statistical Analysis

Statistical analyses were performed for all quantification described above and plotted using GraphPad Prism 9. P values were calculated by performing two-tailed unpaired t-test. In the figures, * = p< 0.05, ** = p< 0.01, *** = p< 0.001, **** = p< 0.0001.

## Supplementary Information

Supplementary Table 1: List of all chemical reagents and antibodies used

Supplementary Table 2: List of smFISH probes used in the study

Supplementary Table 3: Sequences of HCR split-initiator probes used in the study

